# M-current regulates firing mode and spike reliability in a collision detecting neuron

**DOI:** 10.1101/335174

**Authors:** Richard B. Dewell, Fabrizio Gabbiani

## Abstract

All animals must detect impending collisions to escape them, and they must reliably discriminate them from non-threatening stimuli to prevent false alarms. Therefore, it is no surprise that animals have evolved highly selective and sensitive neurons dedicated to such tasks. We examined a well-studied collision detection neuron in the grasshopper *Schistocerca americana* using *in vivo* electrophysiology, pharmacology, and computational modeling. This lobula giant movement detector (LGMD) neuron is excitable by inputs originating from each ommatidia of the compound eye, and it has many intrinsic properties that increase its selectivity to objects approaching on a collision course, including switching between burst and non-burst firing. Here, we demonstrate that the LGMD neuron exhibits a large M current, generated by non-inactivating K^+^ channels, that narrows the window of dendritic integration, regulates a firing mode switch between burst and isolated spiking, increases the precision of spike timing, and increases the reliability of spike propagation to downstream motor centers. By revealing how the M current increases the LGMD’s ability to detect impending collisions our results suggest that it may play an analogous role in other collision detection circuits.

**New & Noteworthy:** The ability to reliably detect impending collisions is a critical survival skill. The nervous systems of many animals have developed dedicated neurons for accomplishing this task. We used a mix of *in vivo* electrophysiology and computational modeling to investigate the role of M potassium channels within one such collision detecting neuron and showed that through regulation of burst firing and increasing spiking reliability the M current increases the ability to detect impending collisions.

## Introduction

Failure to detect an impending collision can have serious, even fatal consequences. So, one might expect that neural circuitry dedicated to this task be highly sensitive. Yet, much of the visual cues of an impending collision are shared by non-threatening stimuli including optic flow, approaching objects on a miss trajectory, or approaching objects slowing to a stop. For this reason, neural circuitry dedicated to detecting impending collisions also needs to also be highly selective.

One of the ways that sensory neurons can be both sensitive and selective is to use multiple firing modes. By multiplexing tonic and burst spiking, neurons increase their ability to both detect and discriminate, or to encode sensory information with both sensitivity and selectivity (Sherman 2001; Krahe and Gabbiani 2004). The use of burst and non-burst spiking as distinct firing modes has been demonstrated in contexts as diverse as auditory neurons in crickets, visual neurons in flies, the electrosensory system of weakly electric fish and the LGN of the mammalian thalamus (Lesica and Stanley 2004; Marsat and Pollack 2006; Longden et al. 2017; Allen and Marsat 2018). In these systems, changes in a neuron’s firing mode signify a change between detection and discrimination. In the detection mode, sensitivity is increased by the generation of a burst of spikes in response to a sudden or novel change in the stimulus. In the discrimination mode, selectivity is increased by encoding stimulus details in the temporal pattern of spikes. The switch between these modes can depend on both intrinsic membrane properties and changes in network activity (Sherman 2001; Krahe and Gabbiani 2004).

Intrinsic neuronal mechanisms involved in switching firing mode include K^+^ conductances, such as the M current (Yue and Yaari 2004; Golomb et al. 2006; Battefeld et al. 2014). The M current, generated by KCNQ/Kv7 channels, is a voltage-dependent, non-inactivating K^+^ current involved in numerous aspects of neuronal function that is evolutionarily conserved with similar properties in mammals, worms, and insects (Wei et al. 2005; Cavaliere and Hodge 2011; Greene and Hoshi 2017). Initial characterizations of the M current showed that a slow depolarization of the postsynaptic membrane potential following activation of muscarinic acetylcholine receptors was due to a deactivation of an M current (Brown and Adams 1980; Adams and Brown 1982). Postsynaptically, the M current contribute to spike frequency adaptation and dendritic integration (Delmas and Brown 2005; Hu et al. 2007; Shah et al. 2011). More recently, a high density of KCNQ channels has been found in the axon and synaptic terminals of various neuron types (Devaux et al. 2004; Vervaeke et al. 2006; Huang and Trussell 2011; Battefeld et al. 2014). These channels contribute to the resting membrane potential (RMP) and diminish excitability (Schwarz et al. 2006; Huang and Trussell 2011; Battefeld et al. 2014).

To explore the neuronal effects of the M-current on the responses of collision detecting circuits, we used an identified neuron in locusts that shows both high sensitivity to small visual objects (Rowell et al. 1977) and selectivity for visual stimuli mimicking impending collision (Schlotterer 1977; Rind and Simmons 1992): the lobula giant movement detector neuron (LGMD; O’Shea and Williams 1974). The LGMD integrates inputs originating from every facet of the compound eye, and despite activation of individual facets reliably triggering spiking (Jones and Gabbiani 2010), it responds selectively to approaching objects that activate thousands of facets based on a simulated object’s trajectory and spatial coherence (Gray et al. 2001; Dewell and Gabbiani 2018). The firing pattern of the LGMD is conveyed to motor centers that initiate and control escape behaviors by a relay neuron the descending contralateral movement detector (DCMD; O’Shea and Williams 1974), and different aspects of its firing pattern including burst firing have been tied to the generation of escape behaviors (Fotowat et al. 2011; McMillan and Gray 2015; Dewell and Gabbiani 2018).

Here we employ this well-characterized neural system with a clear behavioral role to investigate the role of the M-current in the sensory encoding of threatening visual stimuli. Using a mix of *in vivo* electrophysiology, pharmacology, and computational modeling we demonstrate that the M conductance g_M_ narrows the window of dendritic integration by decreasing temporal summation, that it regulates a firing mode switch between burst and isolated spiking, that it increases the precision of spike timing, and that it increases the reliability of spike propagation. Combined, these features are expected to increase the LGMD’s ability to encode the sensory features of approaching objects and help locusts avoid predation.

## Materials and methods

### Animals

All experiments were performed on adult grasshoppers 7-12 weeks of age (*Schistocerca americana*). Animals were reared in a crowded laboratory colony under 12 h light/dark conditions. For experiments, preference was given to larger females ∼3 weeks after final molt that were alert and responsive. Animals were selected for health and size without randomization, and investigators were not blinded to experimental conditions. Sample sizes were not predetermined before experiments. The surgical procedures used have been described previously (Gabbiani and Krapp 2006; Jones and Gabbiani 2012; Dewell and Gabbiani 2018).

### Visual stimuli

Visual stimuli were generated using Matlab and the PsychToolbox (PTB-3) on a personal computer running Windows XP. A conventional cathode ray tube monitor refreshed at 200 frames per second was used for stimulus display (LG Electronics, Seoul, Korea). Looming stimuli are the two-dimensional projection on a screen of an object approaching on a collision course with the animal. They consisted of dark squares simulating the approach of a solid object with half size *l* and constant approach speed *v*, characterized by the ratio *l/|v|* (see top inset in Fig. 3A), as previously described (Gabbiani et al. 2001).

**Figure 1.**
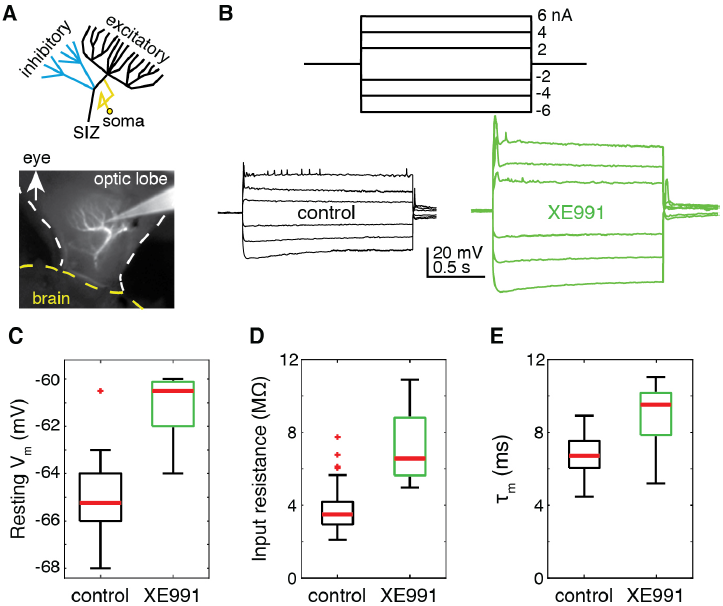
Intracellular LGMD recordings reveal a resting M conductance (g_M_). A) Top, a schematic illustration of the LGMD. The recordings were taken from the region illustrated in black. SIZ, spike initiation zone. Bottom, a micrograph of the LGMD stained with Alexa 594 and the intracellular recording pipette. B) Current steps were injected before (left) and after (right) application of the g_M_ blocker XE991. Both hyperpolarizing and depolarizing currents generated larger changes in membrane potential after g_M_ blockade. Traces have been median filtered to remove spikes. C) The resting membrane potential increased after g_M_ blockade by XE991 (p = 0.0013; control: 11 recordings from 8 animals; XE991: 12 recordings from 8 animals). D) The input resistance increased after g_M_ blockade by XE991 (p = 1.87•10^−7^; control: 79 recordings from 59 animals control; XE991: 13 recordings from 8 animals). E) The membrane time constant (τm) increased after g_M_ blockade by XE991 (p = 4.27•10^−4^; control: 83 recordings from 59 animals; XE911: 13 recordings from 8 animals). In C-E, central red lines are medians, top and bottom box edges are 75^th^ and 25^th^ percentiles, the whiskers denote the extent of data up to 1.5 times the interquartile range and red crosses denote outliers.

**Figure 2.**
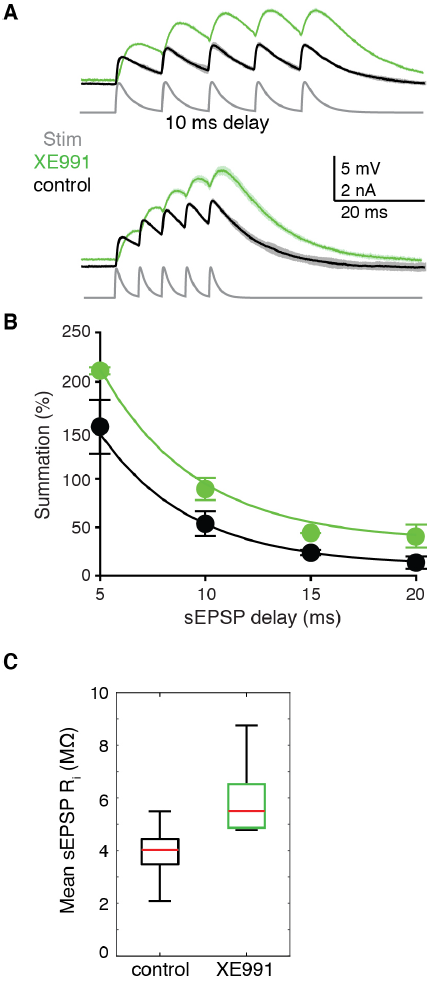
Injection of simulated EPSP currents shows that g_M_ reduces temporal summation. A) Example traces before and after block of g_M_ with XE991. Each trace shows the LGMD membrane potential in response to 5 sEPSP currents (grey traces, Stim) with delays of either 5 ms (bottom) or 10 ms (top). After g_M_ blockade the summation of theses sEPSPs increased for both delays (measured as % change from peak of first to fifth). B) Summation decreased exponentially with longer sEPSP delays. Summation was higher for delays of 5 ms (p = 0.006), 10 ms (p = 0.001), and 20 ms (p = 0.003); data was recorded after XE991 application with a delay of 15 ms for only 1 animal. C) Measuring the mean resistance associated with sEPSP revealed a 50% increase in total sEPSP response after g_M_ blockade by XE991 (p = 3.5•10^−7^). For B and C, control data from 10 recordings from 6 animals and XE991 data from 5 recordings from 3 animals.

**Figure 3.**
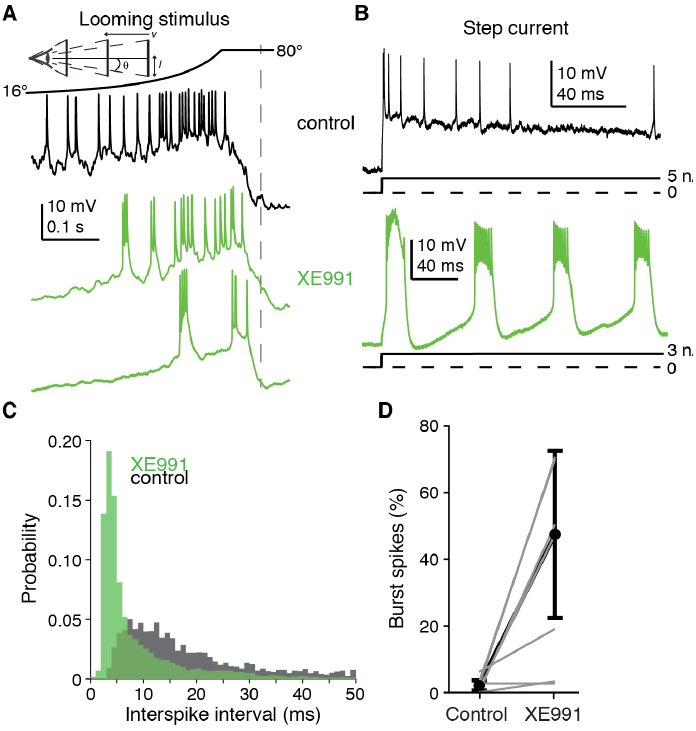
Blockade of g_M_ increased burst firing. A) Example responses to looming stimuli (*l/|v|* = 50 ms) before and after g_M_ blockade by XE991. Top, schematic of looming stimulus, half-size *l*, approach speed *v*, half-angular subtense at the eye,*θ*. Black line shows the non-linear increase in angular subtense (2*θ*), characteristic of looming stimuli. After XE991 addition there was an increase in bursting with clear pauses in firing that were not seen in control looming responses. Dashed line indicates the projected time of collision. B) Depolarizing step currents generated rhythmic bursting after application of XE991 instead of the isolated spikes generated in control.C) The probability histogram of the interspike intervals exhibits a large increase in intervals of ∼4 ms after XE991 addition. D) The prevalence of burst spikes (those with ISIs of 2-5 ms) increased after XE991. Grey lines show data from individual animals (N=8). Black lines show the population response; points are median, error bars are ± mad. Analysis in C and D included 4,684 spikes for control and 14,865 spikes after XE991 application from 8 animals.

### Electrophysiology

Extracellular recordings of the DCMD were carried out with a pair of Formvar coated stainless steel wire hooks placed on the ventral nerve cord between the suboesophageal and prothoracic ganglia (Gabbiani et al. 2001). The DCMD recordings were bandpass filtered from 100 and 5,000 Hz and digitized at 10,036.5 Hz.

Sharp-electrode LGMD intracellular recordings were carried out in current-clamp mode using thin walled borosilicate glass pipettes (outer/inner diameter: 1.2/0.9 mm; WPI, Sarasota, FL; see Jones and Gabbiani 2012, as well as Dewell and Gabbiani 2018 for details). After amplification, intracellular signals were low-pass filtered (cutoff frequency: 10 kHz for the membrane potential, V_m_, and 5 kHz for the membrane current, I_m_) and digitized at a sampling rate of 20,073 Hz. We used a single electrode clamp amplifier capable of operating in discontinuous mode at high switching frequencies (typically ∼25 kHz; SEC-10, NPI, Tamm, Germany). Responses to current injections were recorded in discontinuous current clamp mode (DCC). For two animals we conducted dual recordings (Fig. 4C) by inserting a second sharp electrode into the excitatory dendritic field of the LGMD (Fig. 1A) with a motorized micromanipulator (Sutter Instruments, Novato, CA). Recordings were made in bridge or DCC mode with a second SEC-10 amplifier. Switching frequencies, signal filtering and digitization was the same for both recordings.

**Figure 4.**
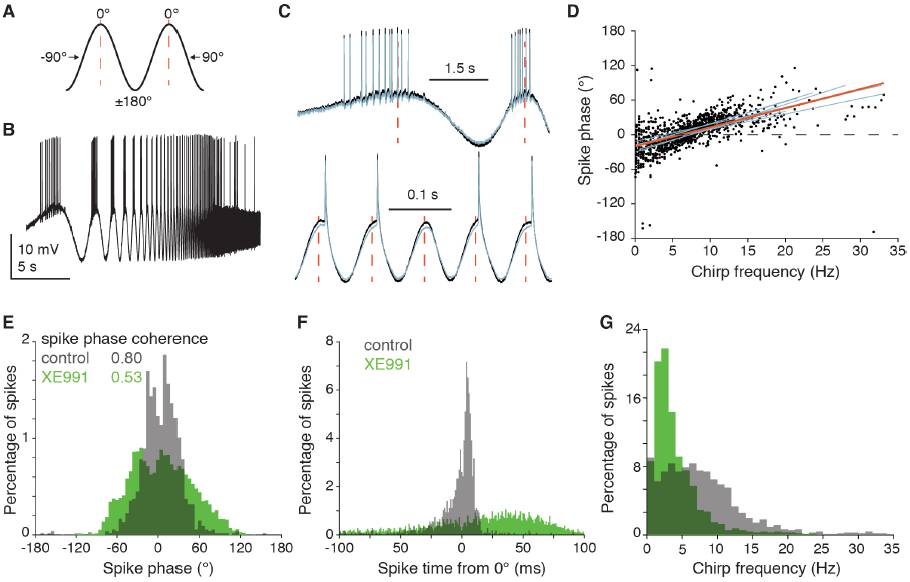
The reliability of spike timing in the LGMD decreased after M channel blockade. A) Illustration of the spike phase measurement. Spikes that occurred at the peaks (dashed red lines) and troughs of the input chirp current were considered to be at 0 and ±180°, respectively. Spikes occurring during the rising slope of the current have a negative phase and those occurring during the falling slope have a positive phase. B) An example trace showing high firing elicited by a chirp current superposed on a depolarizing holding current (2 nA). C) Expanded regions of the same trace as in B plus a simultaneous recording from another dendritic location (grey) show spiking during low (top) and high (bottom) input frequencies. Dashed red lines mark 0° phase. At low input frequencies most spikes are generated on the rising (negative) phase but at high frequencies they come on the falling (positive) phase. The measured spike phase was independent of the recording location (although not necessarily of the current injection location) due to the high synchrony of bAPs across dendritic locations. D) Scatter plot showing the spike phase progression over the chirp current. Linear fits for the population and individual animals are shown in red and blue, respectively, and illustrate the consistency across animals. 0° phase crossing (dashed line) was 6.2±1.2 Hz. 1,074 spikes from 9 animals (r = 0.630 p=5.7•10^−120^). Red line fit, y = a +bx, with a=-20.4° and b=3.33 °/Hz. E) The spike phase probability distribution for control data (gray bars) shows that spikes cluster around 0° with high phase coherence. If the M-current is blocked (green bars; 3,666 spikes from 5 animals) a large reduction in phase coherence is seen (p = 1.2•10^−56^, F-test). Spike phase coherence was >0.65 for all control animals and median spike phase was 5.2°. F) Spike phase was converted to the time domain be dividing by the stimulus frequency (Methods). This revealed that in control conditions spikes were grouped near 0° with 65% within 10 ms of 0° (median ± mad was 1.95±5.94 ms). After XE991 the spike timing was less consistent and more spikes trailed the input current peak (10% within 10 ms of 0° median ± mad was 16.9±38.1 ms). G) For control data there was no clear preferred frequency for spike generation. After blocking the M-current a large proportion of spikes were generated at 2-5 Hz.

**Figure 5.**
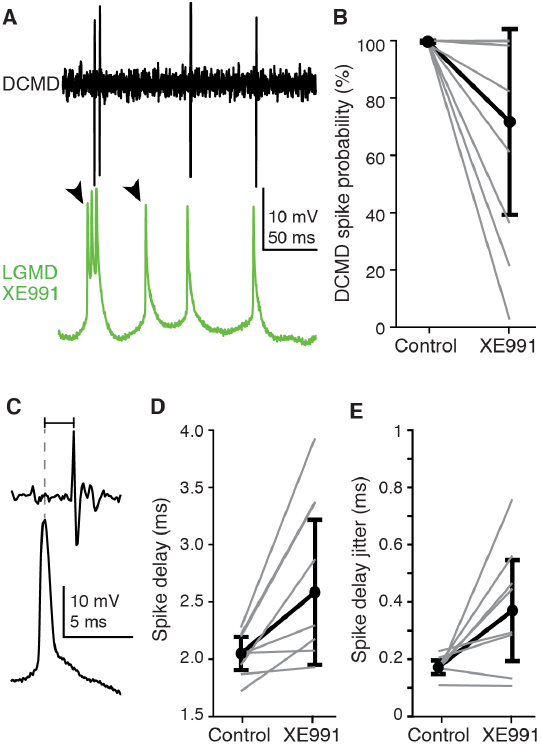
Blocking g_M_ in the LGMD causes failures in DCMD spike initiation. A) Example trace of a LGMD-DCMD recording pair after g_M_ blockade by XE991 (300 mM). Arrowheads mark LGMD spikes that fail to initiate DCMD spikes. B) Under control conditions LGMD spikes reliably initiate DCMD spikes (average of 99.7 % of LGMD spikes had clear matched DCMD spikes with 1-5 ms delay). After g_M_ blockade by XE991 this was substantially reduced. Data from individual animals are displayed with grey lines; 6 of 8 animals had significant reductions in DCMD spike initiations (p < 0.05). On average only 62.7% of LGMD spikes initiated DCMD spikes after g_M_ block; black lines and points are median ± mad of individual percentages. C) For LGMD-DCMD spike pairs the spike delay was measured from the peak of the intracellular LGMD spike (dashed line) to the peak of the extracellular DCMD spike. D) In all 8 animals tested, the spike delay significantly increased after g_M_ blockade with an average increase of 0.7 ms (p<0.01). Individual animal data shown in grey, black lines and points are median ± mad of individual means. E) In 6 of 8 animals there was also a reduction in the consistency of the spike delay (p=0.039, sign rank test) with an average increase in the SD of the spike delay of 0.2 ms (p=0.032, paired t-test).

Injected currents consisted of steps (1-2 s in duration), waveforms producing simulated excitatory postsynaptic potentials (sEPSPs), and chirp currents. sEPSPs were generated by injecting a series of five current waveforms with a set delay between them varied between 5 and 20 ms. Each waveform, *I(t)*, had a time course resembling that of an excitatory postsynaptic current

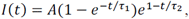

with peak amplitude *A*, rising time constant *τ* _1_ = 0.3 ms, and falling time constant *τ* _2_ = 3.0 ms. This sequence of sEPSPs was used to measure membrane potential summation, calculated as the ratio (p_5_-p_1_)/p_1_, with p_1_ and p_5_ being the peak amplitude of the membrane potential relative to rest during the 1^st^ and 5^th^ sEPSP. We measured the mean sEPSP input resistance by dividing the integrated membrane potential (relative to rest) by the integrated input current (charge) giving a value in units of mV ms / nA ms = MΩ that is readily comparable to the input resistance recorded in response to current pulses.

Chirp currents are sine waves of increasing frequency. They allow rapid and efficient probing of the frequency response of a neuron. If the chirp current is defined as *I*(*t*) =*I*_*p*_sin *ϕ*(*t*), where *I*_*p*_ is the peak current, *ϕ*(*t*) is the phase of the sine wave, then its instantaneous frequency is defined as 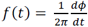 (in units of Hz). Note that if *ϕ*(*t*) = 2*πvt,* the frequency is independent of time and equal to *v*. We used chirps with a duration of 20 s that increased in frequency either linearly or exponentially with time. As the period of a sine wave is inversely related to its frequency, low frequency waves take disproportionally longer to repeat themselves. Hence, using an exponential frequency sweep yielded a more even distribution over time of the frequencies included in the chirp and was used in most experiments, as well as for0020all simulations. The linear chirp started at 0 Hz and was calculated as *I*(*t*) = *I*_*p*_sin *πβt*^2^, with *t* being the time from the start of the chirp (in units of s) and *β* the rate of increase in instantaneous chirp frequency (in Hz/s). The exponential chirp was given by *I(t)* = *I*_*p*_sin2*πf*_0_*te*^*βt*^, where *f*_0_ is the initial chirp frequency and *β* determines the (accelerating) rate of frequency increase (Barrow and Wu 2009). For all exponential chirps, we used *f*_0_ = 0.05 Hz and β = 0.24 Hz which produced a chirp current increasing to 35 Hz over 20 s. To rule out phase delays between the computer-generated waveform and the current injected by the amplifier, we recorded for all chirps the injected membrane current simultaneously with the membrane potential. We used this current for data analysis to prevent any timing delays in the generation of the chirp currents from skewing results. Indeed, over the course of the 20 s chirp duration, we noticed a small timing error(∼0.5 %) originating in the Windows computer generating the chirp currents that could accumulate, causing the end of the current waveform to deviate from the theoretical one by ∼100 ms. Thus, the simultaneous membrane current recordings greatly increased the reliability with which the phase of spikes relative to the oscillating chirp current could be calculated.

### Pharmacology

We applied the M-channel blocker XE991 (10,10-bis (4- pyridinylmethyl)-9(10*H*)-antracenone) either directly in the bath saline or by local puffing. For local puffing, drugs were mixed with physiological saline containing fast green (0.5%) to visually monitor the affected region. They were puffed using a micropipette connected to a pneumatic picopump (PV830, WPI, Saratoga, FL). For bath applications, we used XE991 at concentrations of 200-400 μM and for local puffing we used a concentration of 6 mM in the pipette. Based on previous calculations, this produced concentrations of ∼100–200 μM in the lobula (Dewell and Gabbiani 2018). The LGMD axon travels through the protocerebrum in the brain, with its synaptic terminals located > 500 μm from our puff pipette in the lobula. Our results suggest that M channels extend through the axon, and consequently these channels would be exposed to < 100 μM XE991. In pilot experiments, effects of M current block by XE991 were observed with lobula concentrations <30 μM, but the full effects described in the current experiments required the higher concentration. Previous dosage response measurements have found XE991 to have partial effects at <10 μM, with complete block requiring ∼100 μM (Martire et al. 2004; Cavaliere and Hodge 2011; Battefeld et al. 2014). Thus, the data presented here are likely the result of complete blockade of M–channels within the LGMD. The mechanisms by which XE991 blocks the M current are still under investigation. In isolated cultured cells, 10 μM XE991 was found to only block activated subunits with little effectiveness near RMP (Greene et al. 2017). Here we find that XE991 decreased g_M_ at RMP, as has been found in other neurons (Guan et al. 2011). The difference in these findings might be due to modulatory state of the channels or differences in drug concentration. Although XE991 is selective for M channels, partial block of other K^+^ channels by XE991 has been demonstrated in cultured cells, with 100 μM XE991 reducing K_V_2.1 currents ∼20% in HEK cells (Wladyka and Kunze 2006) and reducing ERG1–2 by ∼20–50% in *Xenopus* oocytes (Elmedyb et al. 2007). It is unknown whether the LGMD possesses these channel types or if they would be affected by these concentrations of XE991 *in vivo*. In one animal, data was collected after washing of XE991 which required >30 min of rinsing with fresh saline. In this preparation, all measurements returned to control levels including 100% DCMD spike propagation and normal spike timing (see Results).

The addition of XE991 reduced excitatory synaptic inputs impinging on the LGMD (see Discussion). To test whether this could explain any of the reported effects, we blocked synaptic inputs with mecamylamine. After mecamylamine application, the RMP was –65.3 ± 1.2 mV, temporal summation was unchanged, burst firing was not increased, and spike timing was not more variable. In each of these cases the effects of mecamylamine were not significant and in the opposite direction of those of XE991. Mecamylamine increased the LGMD input resistance and membrane time constant by ∼5%, but this change is much smaller than that produced by XE991 (Fig. 2D and E).

### Data analysis and statistics

Data analysis was carried out with custom MATLAB code (The MathWorks, Natick, MA). Linear fits were done by minimization of the sum of squared errors. Exponential fits were made with the Matlab function ‘lsqcurvefit’, which minimizes the least squared error between the data and fitting function. Goodness of fit was denoted by R^2^, calculated as one minus the sum of the squared errors of the fit divided by the sum of the squared deviation from the mean of the data. Small hyperpolarizing step currents were used for calculating the input resistance and the membrane time constant. The membrane time constant was calculated by fitting a single exponential to the membrane potential for the period from 0.5 to 13 ms after the start of hyperpolarizing current injection.

Unless otherwise specified, all statistical tests were made using the Wilcoxon rank sum test (WRS) which does not assume normality or equality of variance. For any tests that assumed normality, normality of the data was first assessed by using a Lilliefors test. For displaying summary data, average values were given as median and variance was displayed as median average deviation (mad). For linear fits, the reported ‘r’ and ‘p’ values were calculated from the Pearson linear correlation coefficient testing its statistical difference from zero.

For box plots, the center line shows the median, the upper and lower box limits mark the 25^th^ and 75^th^ percentile of the distribution, and the “whiskers” above and below each box extend 1.5 times the interquartile range up to the minimum and maximum values. Points beyond the whiskers mark outliers.

The timing of spikes was calculated based on their peak for both intracellular and extracellular recordings. During chirp currents, spike phases were calculated relative to the peaks of the sine wave, defined as 0°, with negative phases on the rising portion of the sine wave and positive phases on the falling portion. The throughs of the sine wave corresponded to ±180°. Using Matlab, this was implemented by applying the function ‘acosd’ which computes the inverse cosine in degrees between 0 and 180°, to the normalized chirp current, *I*(*t*)/*I*_*p*_, at the time of a spike, multiplied by minus one times the sign of the first derivative of the chirp current at the time of the spike. The corresponding spike time relative to the time of 0° phase was calculated by dividing the phase by 360° and by the instantaneous chirp current frequency (in s^−1^). The instantaneous chirp frequency was calculated by approximating numerically the first derivative of sin^−1^*I*(*t*)/*I*_*p*_ divided by 2•π. Spike phase coherence was calculated as 1 minus the variance of *e*^iΦ^where 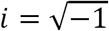 and Φ is the vector of all spike phases (in radians, scaled from -π to π (Sinha and Narayanan 2015). To calculate the 0° phase crossing for each recording, a least squared error linear fit was calculated between the instantaneous chirp frequencies and the spike phases of that recording. The frequency value that produced a spike phase of 0° was computed from the fit line (see Fig. 4D).

To calculate the reliability of the LGMD to DCMD spike propagation we identified DCMD spikes by their peak height from extracellular recordings. For most DCMD recordings, the DCMD spikes are much larger than those of all other neurons in the nerve cord and can be easily distinguished. In noisy nerve cord recordings, some DCMD spikes were not clearly separable from those of other axons in the nerve cord during high frequency firing. In these cases, we were conservative, preferring to exclude DCMD spikes over inclusion of possible non-DCMD spikes. Once DCMD spikes were identified, we iteratively examined each LGMD spike and checked for a matching DCMD spike from 1.3 - 4.8 ms following the LGMD spike. Previous paired DCMD recordings have shown a 1 ms synaptic delay between LGMD and DCMD spikes and ∼1 ms axonal conduction time to the thoracic connective where we record DCMD spikes (O’Shea and Williams 1974; Rind 1984), so we chose the 1.3 - 4.8 ms range to include the expected delay plus a small buffer for any later spikes; in all control spikes the delay fell between 1.6 and 2.5 ms. To prevent a DCMD spike from being matched to 2 LGMD spikes with ISI < 4.8 ms, DCMD spikes were removed upon matching. Even with the conservative thresholding to exclude all possible non-DCMD spikes, a paired DCMD spike was identified for 99.7% of LGMD spikes in control conditions, so a more inclusive analysis would have minimal effects on the findings. The DCMD spike probability was the percent of LGMD spikes with identified matched DCMD spikes. After matching LGMD and DCMD spikes, the spike delay was the time from the peak of the LGMD spike to the peak of the DCMD spike.

### Neuronal modeling

We adapted a recently developed model for the LGMD simulations, described in more detail in Dewell and Gabbiani (2018) and available for download from the ModelDB repository (accession code 195666). For the M conductance, the conductance density varied with neuron location as follows (all conductance values are for −65 mV). The axon had a constant density of 53 μS/cm^2^, the SIZ had a constant density of 144 μS/cm^2^, the primary neurite connecting the dendrites to the SIZ had conductance densities decreasing with distance from the SIZ such that the compartment closest to the SIZ had a density of 226 μS/cm^2^, the compartment at the base of the excitatory dendrites had a density of 79 μS/cm^2^, and the average density was 125 μS/cm^2^. The two inhibitory dendritic subfields had a mean density of 40 μS/cm^2^, and the excitatory dendritic field had a mean density of 17 μS/cm^2^ (see schematics in Fig. 6A). The channels were modeled with steady-state activation set by a Boltzmann function with half activation of −47 mV and steepness of 12 mV, yielding a resting M conductance that was 13.1% of its peak conductance. There were no synaptic inputs or added membrane noise in the simulations. The model and code allowing to reproduce Figures 6 and 7 are available on ModelDB (accession code 241169).

**Figure 6.**
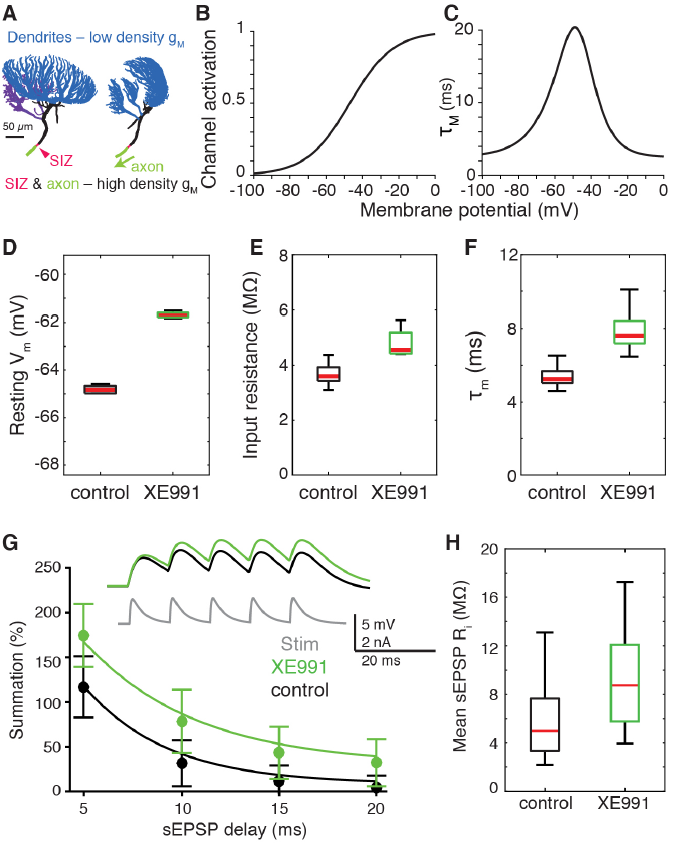
A multi-compartmental model of the LGMD reproduced the subthreshold effects of g_M_.A) Illustration of the model morphology from two orientations. The LGMD’s three dendritic fields (shown in blue) had an average M conductance density of 26 μS/cm^2^ at V_rest_ (−65 mV). The LGMD’s primary neurite (shown in black) which connects the dendritic fields to the SIZ (red) and continues as the axon (green) had an average conductance density of 93 μS/cm^2^ at V_rest_. The axon extends further than pictured and had a total length of 463 μm. B) The g_M_ used for simulations had a broad steady state activation curve with steepness of 12 mV. C) The time constant of g_M_ activation had a minimum and maximum time constants of 2.5 and 21 ms. D) Removal of g_M_ to simulate XE991 application increased the resting membrane potential by 2.7-3.5 mV. For all simulations, data ranges and variability are from measurements of different model sections. E) Measured input resistance to step currents increased by 25-42% after g_M_ reduction. F) Membrane time constant also increased after simulated XE991 application by 41-55%. G) Measured summation to sEPSPs increased for all delays after simulated XE991 application, points and error bars are median ± mad. Inset shows traces of injected current and membrane potential for sEPSPs with delays of 10 ms. H) Median effective input resistance to sEPSPs increased from 5.0 to 8.7 MΩ after g_M_ blockade.

**Figure 7.**
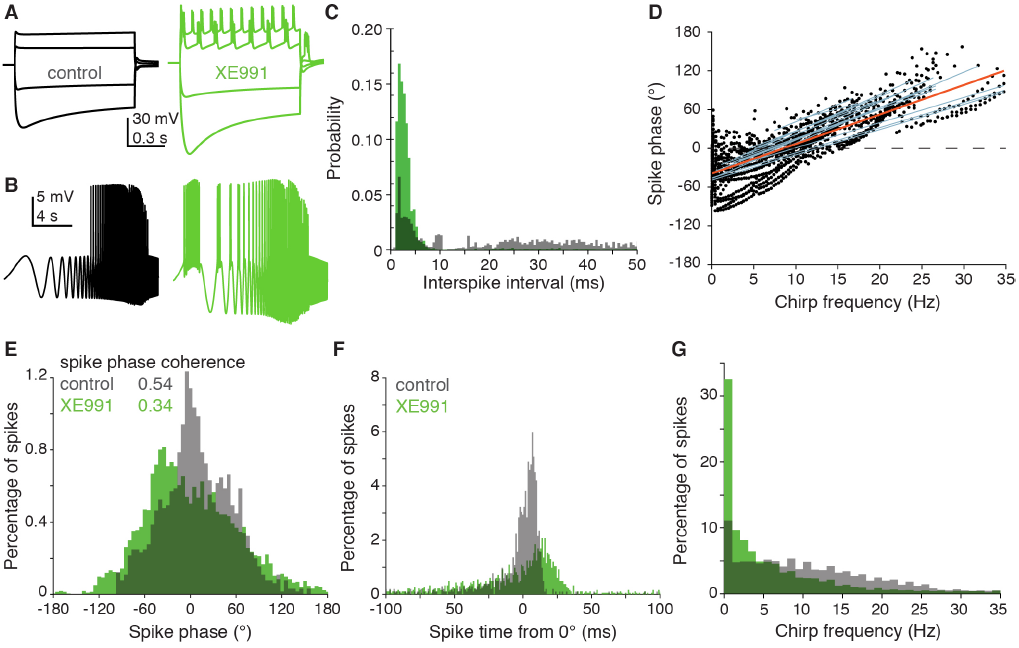
Model LGMD reproduced the role of g_M_ in the LGMD’s spiking pattern. A, B) Simulated effect of g_M_ blockade on depolarizing current steps and chirp currents injected into different dendritic sections to induce spiking. For control chirp currents, a constant depolarizing current was superposed (2.5 nA in example trace). C) After g_M_ blockade, there was a shift in the ISI distribution with the proportion of spikes with ISIs of 2-5 ms increasing from 13 to 44%. D) As in experimental data, a spike phase progression occurred with increasing chirp frequency. Data displayed as in Fig. 4D but with each line representing data recorded from different model compartments instead of different recordings. E) A broadening of the spike phase distribution produced a 37% reduction in spike phase coherence after g_M_ blockade. F) The decreased reliability in spike time is seen after converting the phase to time from peak current. The spikes occurring within 10 ms of 0° decreased from 62 to 19 % after g_M_ blockade (compare with Fig. 4F). G) The model LGMD also produced a shift to a larger percentage of spikes being generated by low frequency inputs.

## Results

### g_M_ alters the electrotonic properties of the LGMD

To test for the presence and role of g_M_ in the LGMD, sharp electrode intracellular recordings were employed *in vivo* before and after application of the specific g_M_ blocker XE991 (Fig. 1A). After blockade, immediate changes were seen in the passive electrotonic properties of the neuron revealing both that M channels are present within the LGMD and that a significant number are open at rest. This was clear in the increased membrane potential change in response to both hyperpolarizing and depolarizing step currents (Fig. 1B). The resting membrane potential increased by ∼4 mV from −65 to −61mV after XE991 application (Fig. 1C). This increase brings the neuron closer to spike threshold and increased excitability. The median membrane input resistance increased by 88% after g_M_ blockade (Fig. 1D). A single exponential function was fit to the membrane potential for the time period immediately following the onset of each current step to measure the membrane time constant (as in Dewell and Gabbiani 2018). After g_M_ blockade, the median membrane time constant increased by 44% (Fig. 1E). Each of these changes to the electrotonic properties of the membrane after XE991 application suggest excitatory effects consistent with the removal of a K^+^ current.

### g_M_ decreases summation of excitatory postsynaptic potentials (EPSPs) in the LGMD

The conductance g_M_ can reduce EPSP amplitude and summation thereby lessening dendritic integration, as is the case, e.g., in cortical neurons. To test if that is also the case in the LGMD, we injected brief, transient currents to simulate a series of 5 EPSPs at the level of the LGMD’s membrane potential. As shown previously, these simulated EPSPs (sEPSPs) summate sublinearly in the dendrites of the LGMD (Dewell and Gabbiani 2018). Blocking g_M_ with XE991 increased the amplitude and duration of the sEPSPs (Fig. 2A). In addition, the summation from the first to fifth sEPSP increased after g_M_ block by an average of 75% (Fig. 2B). Dividing the mean sEPSP amplitude by the mean current injected yielded the effective input resistance to the sEPSP. This input resistance increased after g_M_ block (Fig. 2C) by an amount similar to that observed during current injection (c.f., Fig. 1D). The increase in sEPSP amplitude and summation after g_M_ block shows that g_M_ reduces excitability to excitatory synaptic inputs.

### g_M_ toggles the firing mode of the LGMD

KCNQ channels can also alter the firing properties of a neuron, so we examined the influence of g_M_ on the spiking output of the LGMD and found dramatic changes. In response to objects approaching on a collision course or their two-dimensional projections, the LGMD generates a characteristic firing pattern. The spike rate ramps up as the stimulus expands, reaches a peak and decays before the projected time of collision. An example of the LGMD response to a looming stimulus is shown in Fig. 3A. In the control case, the usual response was observed. However, after XE991 application, the LGMD responded with bursts of activity, as seen in examples from two animals (Fig. 3A). The increase in burst firing was even more striking for depolarizing current steps. In control conditions such current steps generated immediate firing with a rate that decayed exponentially (Fig. 3B, top; Gabbiani and Krapp 2006; Peron and Gabbiani 2009). After XE991 addition however, the LGMD often generated rhythmic bursts of activity (Fig. 3B, bottom). Examination of the probability histogram of the interspike intervals (ISIs) showed a clear increase in spikes with ISIs around 4 ms (Fig. 3C). In 5 of 8 animals an increase in bursting was seen (p < 0.001), and on average the fraction of spikes having ISIs of 2-5 ms increased from 2% to 38% (Fig. 3D).

### g_M_ improves the reliability of LGMD spiking

A shift in firing mode from isolated to burst spiking has been associated with a change in a neuron’s operating regime from fine discrimination to detection. In the detection mode, whether or not spikes are present conveys the relevant information, while in the discrimination mode specific stimulus parameters may be encoded in the temporal pattern of the spikes. How much information can be conveyed in the timing of spikes depends on their reliability. To test this reliability in the LGMD we injected chirp currents (sine waves of increasing frequency, Methods) at depolarized holding potentials and measured the phase of the generated spikes (Fig. 4A, B). For these measurements, spikes occurring at the peak and the trough of the input current were assigned phases of 0 and ±180°, respectively. Negative and positive phases were assigned to spikes occurring on the rising and falling slope of the sinusoids, respectively (Fig. 4A). For low frequency oscillating currents, spikes mostly came on the rising slope while they fell on the falling slopes at high frequency (Fig. 4C). The measured spike phase was independent of recording location, as shown by the simultaneously recorded and overlaid traces in Fig. 4C. Hence, there was no noticeable propagation lag between simultaneously recorded locations across the excitatory dendritic field. The spike phase was consistent across animals (blue lines) and showed a clear progression with the input frequency of the chirp (Fig. 4D). On average, spikes were most synchronous with the input around theta frequency (zero crossing of blue lines in Fig. 4D). The consistency of spike timing was reflected in their spike phase coherence, that was high compared to other systems (Fig. 4E, grey bars; van Brederode and Berger 2008; McLelland and Paulsen 2009). The spike phase coherence was even higher within narrow frequency ranges, e.g. spikes during the 5-10 Hz period of the chirp had a spike phase coherence of 0.93. After addition of XE991, there was a reduction in spike phase coherence (Fig. 4E, green bars). To compare the absolute difference in spike timing, we plotted the time of each spike relative to the peak of the input current sinusoid (0°). In control data these spike times were tightly grouped around the sinusoid peak time with 65% of spikes occurring within ±10 ms of it (Fig. 4E, grey bars). After XE991 application only 10% of spikes occurred in the same interval (Fig. 4E, green bars). This demonstrates that g_M_ increases the reliability of the LGMD’s spike timing.

### g_M_ is critical for signal propagation from the LGMD to the DCMD

The block of g_M_ not only reduced the spike timing reliability within the LGMD, but also the reliability of the signal propagation from the LGMD to its downstream target the DCMD. Under control conditions, LGMD spikes faithfully generate spikes in the DCMD through a mixed chemical and electrical synapse (Rind 1984; Killmann and Schürmann 1985). We used simultaneous intracellular LGMD and extracellular DCMD recordings to measure how consistent this spike propagation was before and after g_M_ blockade. After XE991 application spike failures occurred during both burst and isolated spiking (Fig. 5A). For control recordings, we detected matching DCMD spikes for 99.7% of LGMD spikes. Manual inspection of the recordings suggested that the 0.3% failure rate was due to limitations in detecting DCMD spikes from noisy extracellular recordings rather than genuine failure of the LGMD/DCMD synapse. Following g_M_ blockade, 6 of the 8 animals had a significant increase in failures of spike transmission to the DCMD, with only 63% of LGMD spikes initiating DCMD spikes on average (Fig. 5B). During control conditions, reliable DCMD spike initiation occurs over a broad range of instantaneous LGMD firing frequencies (0–500 spk/s), so the increase in firing rates after XE991 application cannot explain the failures.

We next measured the time lag from the peak of each LGMD action potential to the peak of the corresponding extracellular DCMD spike (Fig. 5C). In control conditions DCMD spikes trailed those of the LGMD by ∼2 ms. After XE991 application there was a significant increase in this delay in all animals (Fig. 5D). Not only did the delay increase, but its variability increased as well, approximately doubling from 0.17 to 0.37 ms (Fig. 5E). This shows that g_M_ increases the reliability of LGMD spike generation (Fig. 4E, F) as well the reliability of their propagation to downstream motor centers.

### A multi-compartmental model reproduced the subthreshold effects of g_M_

We adapted a recent biophysical model of the LGMD, illustrated in Fig. 6A to investigate if a g_M_ conductance could account for the experimental findings described above. In the model, g_M_ was highest within the axon, the spike initiation zone (SIZ), and the primary neurite connecting the dendritic fields to the SIZ. The dendrites had a lower g_M_ density. Although the exact M-channel kinetics within the LGMD is uncharacterized, the increased input resistance after g_M_ block at both depolarized and hyperpolarized potentials (Fig. 1B, D) suggest a broad activation range. Accordingly, we chose activation kinetics with a shallow slope (Fig. 6B). Simulation with different kinetics than those in Figure 6 were conducted in preliminary simulations. While most properties described were reproducible in kind by a wide range of channel kinetics, faster channel activation (Fig. 6C) improved the similarity between experimental and simulation data. In these simulations removal of g_M_ reproduced the increase in resting membrane potential (Fig. 6D), input resistance (Fig. 6E), and membrane time constant (Fig. 6F) found in experiments after g_M_ blockade by XE991. We simulated current injections with the same time course as the sEPSPs injected experimentally (Fig. 2) and the model also showed increased summation following g_M_ blockade (Fig. 6G). The increased summation after g_M_ blockade was paired with an increased mean input resistance for the sEPSPs (Fig. 6H).

### Dependence of LGMD spiking mode on g_M_ reproduced by LGMD model

We simulated the same currents used in experiments to induce spiking. As in experimental data, depolarizing current steps produced rhythmic bursting after g_M_ removal (Fig. 7A). Similar bursts were also produced by chirp currents (Fig. 7B). These increased bursts produced a shift in the ISI distribution after g_M_ blockade with a larger proportion of spikes having short ISIs (Fig. 7C). The model also exhibited a spike phase progression during the chirp currents with most spikes being produced during the rising phase of the input sinusoid at low frequencies but coming on the falling phase at high chirp frequencies (Fig. 7D). Surprisingly, the spread in the data was higher across locations within the model than it was across animals in our experiments despite the lack of noise sources in the simulations. The LGMD had a spike synchrony frequency of 6.2 ± 1.2 Hz (mean ± SD) measured across multiple animals, which for the model simulations was 6.7 ± 3.8 Hz measured across dendritic locations (Fig. 7D, blue lines). This increased spread of spike phases resulted in a broader spike phase histogram and lower spike phase coherence in simulations than in experimental data (Fig. 7E). In both model and experiment, g_M_ blockade decreased the reliability of the spike phase and lowered spike phase coherence (Fig. 7E; c.f. Fig. 4E). As for spike phase, the time of spiking from the peak of the input current sinusoid was smaller before g_M_ block (Fig. 7F; c.f. Fig. 4F). That removal of g_M_ within the model was sufficient to reproduce the effects of LGMD firing in kind supports the hypothesis that the changes in LGMD firing patterns observed experimentally are primarily intrinsic to the LGMD and not a network phenomenon.

## Discussion

M/Kv7/KCNQ channels influenced both sub- and supra-threshold activity within the LGMD. They increased the reliability of propagation of LGMD activity to downstream motor centers that control escape behaviors, failures of which could have dire consequences for the animal’s ability to escape predation. The resting g_M_ lowered input resistance, membrane potential, and membrane time constant. This led to decreased amplitude and temporal summation of dendritic inputs which would narrow the temporal window of integration. Additionally, blockade of g_M_ caused a change in firing mode greatly increasing the proportion of burst firing, and decreased spike timing consistency. A single cell model replicated all effects in kind if not in exact detail (other than the spike propagation which would require a multi-cell model) confirming the hypothesis that these are direct changes of the LGMD’s intrinsic properties and not indirectly caused by changes in network activity.

Despite combining data from 9 animals, obtained in recordings at 12 different dendritic locations, that caused differing amounts of spiking from chirps in response to different current amplitudes, the population spike phase coherence was high: 0.93 for input frequencies of 5-10 Hz and 0.80 for input frequencies of 0-35 Hz. This is in contrast to pyramidal neurons, where spike phase often changes with recording location, current amplitude, and spike count, a feature believed to play a role in theta spike phase precession (Magee 2001; Harris et al. 2002; McLelland and Paulsen 2009). During looming stimuli, the LGMD receives a broad range of input frequencies and amplitudes. Yet, it generates a firing profile with a consistent time course, independent of the amount of spiking. The consistency of its spiking phase independent of the location or amplitude of the input current might thus help the LGMD generate its characteristic looming response profile. Under control conditions, the spike pattern was consistently relayed to the DCMD (Fig. 5E) allowing this spike pattern to be accurately conveyed to motor centers.

Somewhat surprisingly, blockade of M channels can result in either an increase or decrease in synaptic release (Vervaeke et al. 2006; Shah et al. 2008; Huang and Trussell 2011). These seemingly contradictory effects are believed to be due to an increase in RMP both bringing the membrane closer to spike threshold and increasing the resting inactivation of Na^+^ channels in the axon and of Ca^2+^ channels in the synaptic terminals (Vervaeke et al. 2006; Battefeld et al. 2014; Greene and Hoshi 2017). These competing excitatory and inhibitory effects can cause a simultaneous increase in excitability and decrease in synaptic release. Similar effects most likely cause the LGMD-DCMD spike failures (Fig. 5A, B). A prolonged increase in RMP in the axon could prevent the Na^+^ channels from sufficiently de-inactivating to conduct spikes, or a prolonged inactivation of Ca^2+^channels might have caused a decrease in synaptic release.

The initial characterization of the M current was through its suppression by muscarinic acetylcholine receptor activation (Adams and Brown 1982; Delmas and Brown 2005). Since then, many more modulators of M channels have been discovered. The M current can also be suppressed through activation of 5-HT, substance P, glutamate, opioid, and angiotensin receptors (Greene and Hoshi 2017). Conversely it can be augmented by somatostatin, corticostatin, and dynorphin receptor activation (Greene and Hoshi 2017). These modulators involve numerous different pathways and messengers including PIP_2_, IP_3_, PLC, PKC, Ca^2+^, cyclic nucleotides, and tyrosine kinase (Delmas and Brown 2005). The array of modulation effectors suggests that the LGMD’s firing mode and spike timing might be dynamically regulated by M channel modulation. While we don’t know which, if any, of these modulatory pathways are used to modulate M channels within the LGMD, conducting all experiments *in vivo* ensures that the modulation of the channels during our experiments was in a relevant state.

The presynaptic terminals of medullary excitatory inputs to the LGMD possess mAChRs whose activation leads to lateral excitation, although it is unknown whether activation of these receptors suppresses a presynaptic M current (Rind and Leitinger 2000; Zhu et al. 2018). In our experiments, application of XE991 reduced excitatory synaptic inputs to the LGMD in all animals even with local puffing onto the LGMD. This reduced excitatory input cannot have produced any of the effects described in the results (see Methods). This suggests that there is an M-current in presynaptic terminals or axons of these medullary inputs and that its blockade by XE991 causes a reduction in synaptic release. Activation of mAChRs has an excitatory effect on synaptic release (Zhu et al. 2018) and, as shown here, XE991 has an inhibitory effect on release from medullary inputs to the LGMD. However, this last result does not exclude that the mAChR related excitation is mediated by M current suppression because XE991 drug application might also cause an increase in the resting inactivation of Na^+^ and Ca^2+^ channels leading to the decreased transmitter release (Schwarz et al. 2006; Huang and Trussell 2011; Battefeld et al. 2014). Indeed, this is likely the cause for the spike propagation failures shown in Figure 5.

The LGMD has a large dynamic range that allows strong responses to small visual stimuli (Rowell et al. 1977; Jones and Gabbiani 2010), while still selecting between inputs that activate thousands of facets (Gray et al. 2001; Peron and Gabbiani 2009; Dewell and Gabbiani 2018). This is accomplished through a combination of network and intrinsic properties, with the intrinsic properties including active membrane conductances in the dendrites and at the spike initiation zone that filter synaptic inputs (Gabbiani and Krapp 2006; Peron and Gabbiani 2009; Jones and Gabbiani 2010; Dewell and Gabbiani 2018) and regulation of the spike frequency profile and burst spiking (Gabbiani et al. 2001; Fotowat and Gabbiani 2007; McMillan and Gray 2015; Dewell and Gabbiani 2018). The wide dynamic range of the LGMD helps it detect impending danger and collisions both sensitively and selectively. In many neurons, the M current increases the output range through regulation of excitability, spike frequency, and burst firing (Yue and Yaari 2004; Gu et al. 2005; Lawrence et al. 2006; Schwarz et al. 2006; Vervaeke et al. 2006; Hu et al. 2007; Shah et al. 2008; Battefeld et al. 2014). Here, for the first time to the best of our knowledge, we investigated the role of the M current in a collision detecting neuron. We found that the M current regulates the LGMD’s bursting and spike timing, and that it increases the reliability of the signal propagation from the LGMD to downstream motor centers. These results suggest that neuromodulation through M channels may play a similar role in other collision detection circuits as well.

## Acknowledgements

We would like to Dr. Ed Cooper for feedback on a previous version of this manuscript.

